# Nucleus reuniens is organised as parallel circuits rather than a prefrontal-hippocampal relay

**DOI:** 10.64898/2026.08.11.744129

**Authors:** Jessica Passlack, Kristina Valentinova, Andrew F. MacAskill

## Abstract

Interactions between the prefrontal cortex (PFC) and hippocampus (HPC) are crucial for flexible and memory guided behaviour. Because there are no direct projections from PFC to HPC, it is widely presumed that information is shared through a disynaptic projection via the nucleus reuniens (nRE), a ventral midline thalamic nucleus. Yet this model relies on the implicit assumption that has not been directly tested: that prefrontal afferents monosynaptically recruit hippocampal-projecting nRE neurons. Here we combine whole-brain anatomical mapping, projection-specific circuit tracing and electrophysiology to test this model. We find that nRE contains largely distinct populations of prefrontal- and hippocampal-projecting neurons, with minimal overlap between them. Unexpectedly, prefrontal afferent input to nRE provides little anatomical or functional input to hippocampal-projecting neurons, instead preferentially targeting neurons projecting back to PFC. These findings challenge the prevailing view that nRE acts as a simple relay between PFC and HPC and instead reveal parallel projection-defined circuits likely to make distinct contributions to hippocampal-prefrontal communication.

## INTRODUCTION

Functional interaction between the hippocampus (HPC) and prefrontal cortex (PFC) is thought to underlie flexible behaviour by supporting learning, memory and decision making^1–10^. Activity in these regions becomes synchronised during contextual learning and inference^11–17^, and disrupting either structure produces remarkably similar behavioural deficits^11,18–21^. Consequently, communication between HPC and PFC is widely considered a fundamental mechanism by which contextual information guides behaviour^7,9^.

Despite this, the anatomical basis of this interaction remains surprisingly poorly understood. Whereas the hippocampus sends direct projections to the prefrontal cortex^22,23^, there is no corresponding direct projection from PFC back to HPC^24^. Instead, the thalamic nucleus reuniens (nRE) has emerged as the putative bidirectional link between these regions. nRE receives input from both HPC and PFC^25–27^ and projects back to both structures^28–30^, leading to the prevailing view that it acts as the principal relay through which information is exchanged between them^31,32^. In particular, because no direct PFC to HPC projection exists, many current models assume that nRE provides the pathway by which prefrontal activity influences hippocampal processing during flexible behaviour^13,17,31,33–37^.

However, this assumption has not been directly tested. While previous anatomical studies have established that nRE projects to both HPC and PFC, it remains unclear whether the same neurons innervate both structures, whether the two projection populations receive similar afferent inputs, or indeed whether hippocampal-projecting nRE neurons receive substantial input from PFC. These questions are fundamental because the organisation of the underlying circuit determines which patterns of information flow are biologically possible, and therefore constrains mechanistic interpretations of behavioural experiments.

Here we combine quantitative whole-brain anatomy with projection-specific transsynaptic tracing and circuit mapping to define the organisation of the HPC-PFC circuit through nRE. We find that nRE contains largely distinct populations of neurons projecting either to HPC or to PFC, with very little overlap between them. These projection-defined populations receive distinct afferent inputs and, unexpectedly, hippocampal-projecting nRE neurons receive remarkably little direct input from PFC, a finding confirmed using projection-specific electrophysiological circuit mapping. Together, our results challenge the prevailing view of nRE as a simple bidirectional relay between HPC and PFC and instead reveal parallel projection-defined circuits that are likely to make distinct contributions to hippocampal-prefrontal communication.

## RESULTS

### nRE exhibits reciprocal connectivity with both prefrontal cortex and hippocampus

The nucleus reuniens (nRE) has been proposed to mediate communication between prefrontal cortex (PFC) and hippocampus (HPC), but the extent of its reciprocal connectivity with these regions has not been quantified systematically. We therefore first sought to define the major inputs and outputs of nRE within a common anatomical framework. To identify afferent inputs, we injected cholera toxin B (CTB) into nRE and registered labelled neurons to the Allen Common Coordinate Framework (Fig. 1A–C, Figure S1).

**Figure 1.**
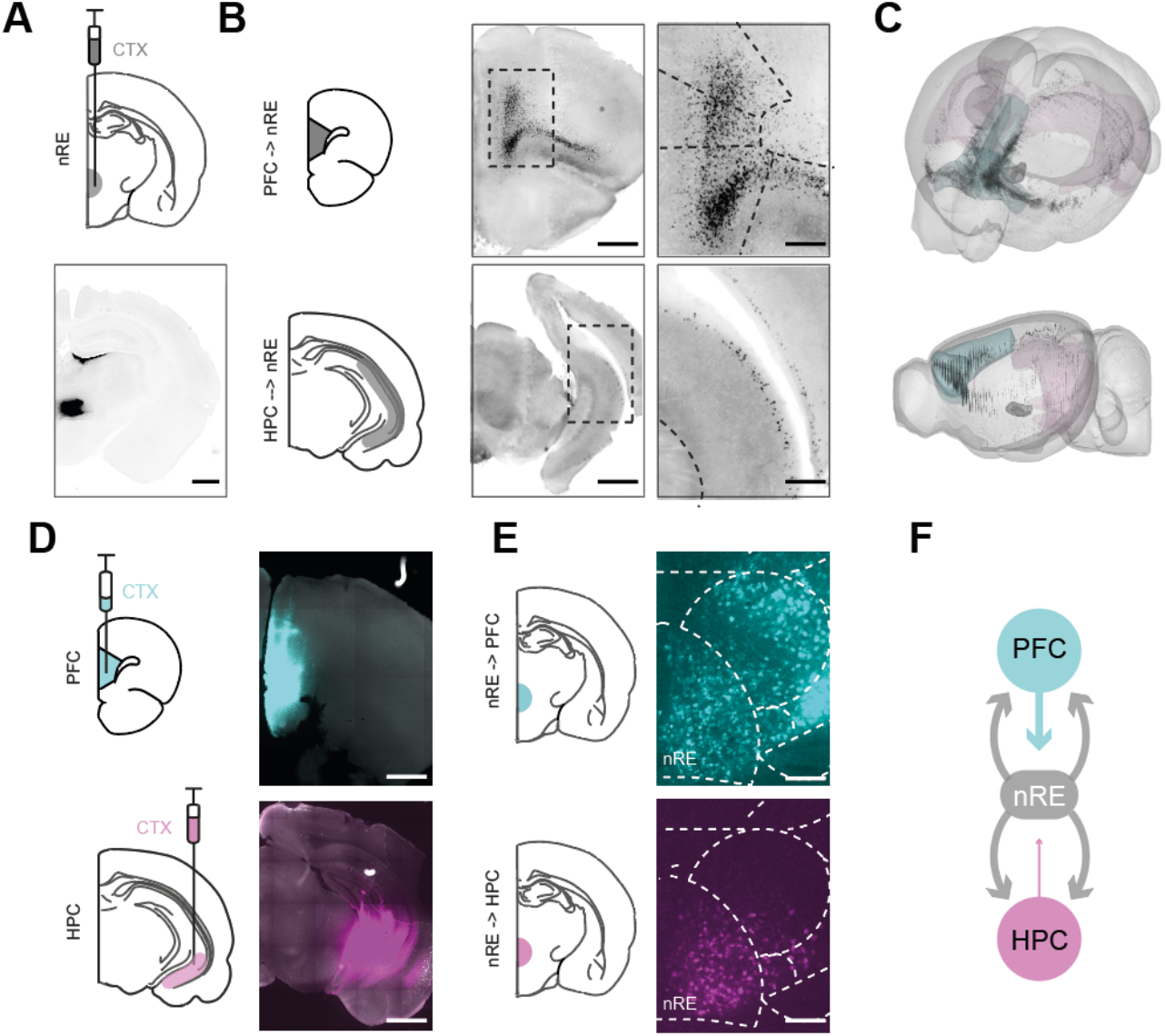
Nucleus reuniens exhibits reciprocal connectivity with both prefrontal cortex and hippocampus. (A) Top, schematic illustrating cholera toxin B (CTB) injections into the nucleus reuniens (nRE) to identify afferent inputs. Bottom, representative CTB injection site in nRE. Scale bar, 1 mm. (B) Representative examples of CTB-labelled neurons following nRE injections in medial prefrontal cortex (PFC; top) and hippocampal formation (HPC; bottom). Scale bars 1 mm (left), 200 μm (right). (C) Whole-brain registration of CTB-labelled neurons following nRE injections, revealing widespread afferent inputs with prominent labelling in PFC and HPC. (D) Schematic (left) and representative injection sites (right) for CTB injections into medial PFC (blue) and hippocampal formation (purple) used to identify nRE projection neurons. (E) Representative thalamic sections following CTB injections into medial PFC (top) or hippocampal formation (bottom), showing dense retrograde labelling within nRE. Scale bars 1 mm. (F) Summary schematic of reciprocal connectivity between nRE, medial PFC and hippocampal formation identified by retrograde tracing.

Following CTB injections into nRE, labelled neurons were distributed throughout the brain but were concentrated within isocortex and hippocampal formation (Fig. 1B,C; Figure S1). Within cortex, the strongest inputs arose from medial and orbital prefrontal regions, including classic medial prefrontal cortical regions such as prelimbic, infralimbic, and anterior cingulate cortex, whereas within the hippocampal formation inputs originated from subiculum, CA1 (mainly in ventral areas) as well as from lateral entorhinal cortex (Fig. 1B,C; Figure S1), but these were markedly fewer than the dense medial prefrontal inputs. Thus, consistent with classical observations nRE receives dense input from mPFC, as well as a smaller input from ventral CA1 and subiculum.

Having established the principal afferent inputs to nRE, we next asked whether these connections were organised reciprocally. We therefore injected CTB into medial PFC or hippocampal formation to identify the major nRE outputs. CTB injections into medial PFC revealed that nRE was one of the major thalamic sources of input to PFC, alongside local cortical projections (Fig. 1D,E; Figure S2). Similarly, injections into ventral CA1 and subiculum identified nRE as the dominant thalamic input to the hippocampal formation (Fig. 1D,E; Figure S3).

Together, these experiments demonstrate that nRE is reciprocally connected with both medial prefrontal cortex and hippocampal formation, confirming its anatomical position as a major hub linking these structures.

### Prefrontal cortex has limited connectivity with hippocampal-projecting nRE neurons

As nRE projects to both PFC and HPC, these outputs could arise either from collateralising neurons or from distinct projection-defined populations. We therefore next asked how PFC- and HPC-projecting neurons are organised within nRE.

To address this, we used dual retrograde tracing from PFC and HPC in the same animals. This revealed minimal overlap between PFC- and HPC-projecting nRE neurons, indicating that these projections arise predominantly from separate neuronal populations (only 2% of nRE cells projected to both regions, Fig. 2A,B).

**Figure 2.**
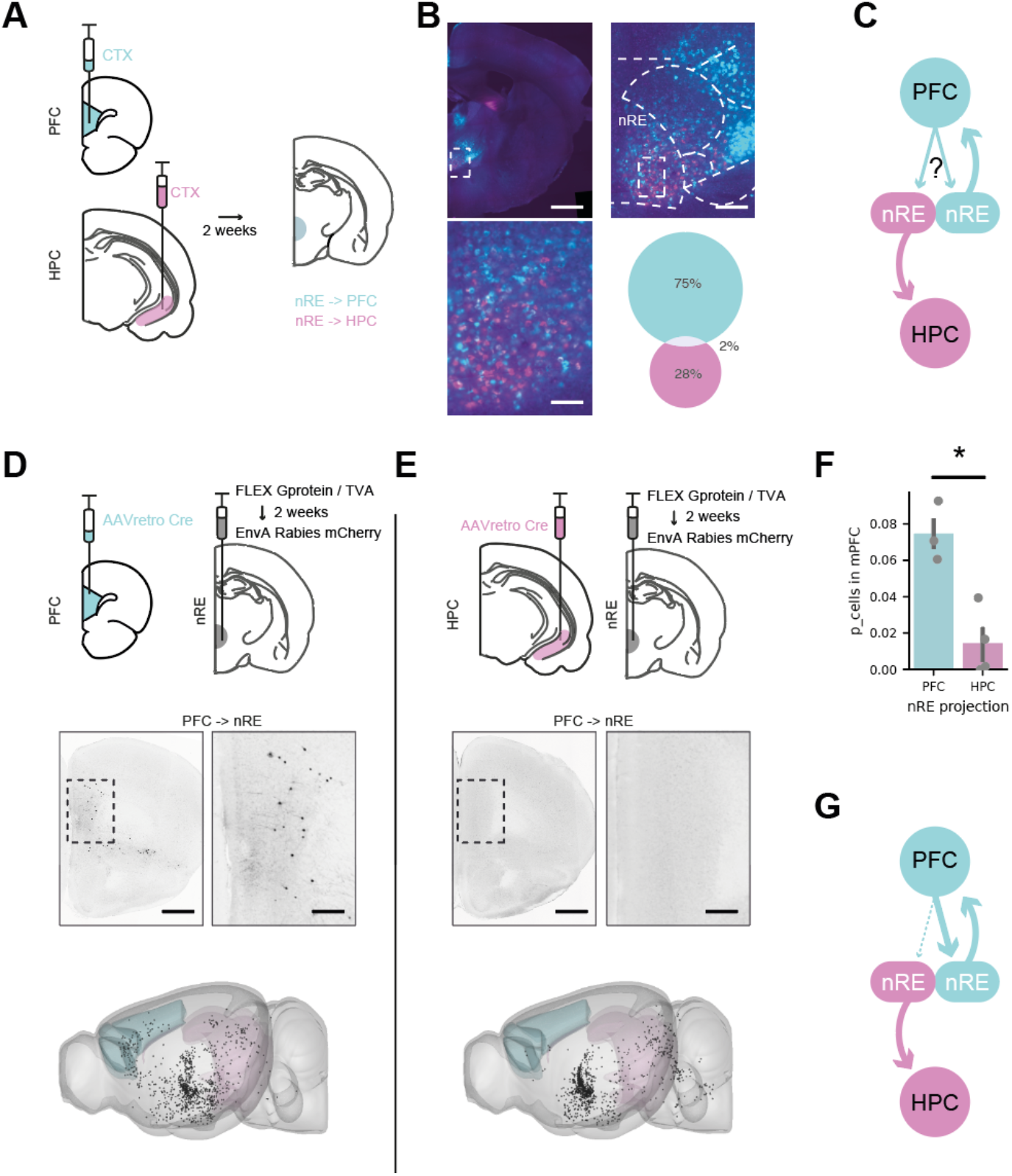
Prefrontal cortex provides little direct input to hippocampal-projecting nucleus reuniens neurons. (A) Schematic illustrating dual cholera toxin B (CTB) injections into medial prefrontal cortex (PFC; blue) and hippocampal formation (HPC; purple) to identify projection-defined nRE neurons. (B) Representative sections through nRE showing retrogradely labelled PFC-projecting (blue) and HPC-projecting (purple) neurons. Dual-labelled neurons were rare. Right, quantification of overlap between projection-defined populations. Scale bars 1 mm, 200 μm, 50 μm. (C) Schematic illustrating the question addressed by projection-specific input mapping: whether prefrontal afferents innervate both PFC- and HPC-projecting nRE neurons. (D) Top, strategy for tracing monosynaptic inputs to PFC-projecting nRE neurons using TRIO. AAVretro-Cre was injected into PFC, Cre-dependent TVA and rabies glycoprotein were expressed in nRE, and EnvA-pseudotyped rabies virus was subsequently injected into nRE. Middle, representative medial PFC section showing rabies-labelled presynaptic neurons. Bottom, whole-brain registration of labelled inputs from a representative animal. Scale bars 1 mm, 200 μm. (E) TRIO strategy for tracing monosynaptic inputs to HPC-projecting nRE neurons. Top, experimental schematic. Middle, representative medial PFC section showing sparse rabies labelling. Bottom, whole-brain registration of labelled inputs from a representative animal. (F) Quantification of rabies-labelled neurons in medial PFC across animals for PFC- and HPC-projecting nRE populations. (G) Summary schematic illustrating the organisation of nRE circuitry. PFC- and HPC-projecting neurons comprise largely distinct populations, and prefrontal afferents preferentially target PFC-projecting rather than HPC-projecting neurons

This anatomical organisation has important implications for prevailing models of nRE function. If PFC- and HPC-projecting neurons were connected locally within nRE, or if individual neurons collateralised to both targets, prefrontal activity could readily be transmitted to hippocampus through the proposed relay. However, previous studies have shown that neurons within nRE lack local recurrent connectivity^38–41^, and our dual tracing experiments demonstrate that PFC- and HPC-projecting neurons are largely distinct. Together, these observations indicate that the canonical PFC–nRE–HPC relay can operate only if prefrontal afferents directly innervate hippocampal-projecting nRE neurons.

We therefore used *tracing the relationship between inputs and outputs* (TRIO) to determine the monosynaptic inputs to projection-defined nRE neurons. To selectively label presynaptic inputs to either PFC- or HPC-projecting nRE neurons, we injected AAVretro-Cre into the projection target, expressed Cre-dependent TVA receptor and rabies glycoprotein in nRE, and subsequently injected EnvA-pseudotyped rabies virus into nRE. This strategy selectively labels neurons providing direct input to the chosen projection-defined population (Fig. 2D,E).

In both experiments, rabies-labelled neurons were distributed throughout the brain, consistent with the diverse afferent connectivity of nRE (Figure S4). However, striking differences emerged within frontal cortex. PFC-projecting nRE neurons received extensive input from classic medial prefrontal regions, including anterior cingulate, prelimbic and infralimbic cortex (Fig. 2D). In contrast, medial prefrontal input to HPC-projecting neurons was almost completely absent, with only sparse labelling observed in neighbouring frontal regions such as orbitofrontal cortex (Fig. 2E). Quantification of rabies-labelled neurons across the major medial prefrontal regions (anterior cingulate, prelimbic and infralimbic cortex) confirmed that HPC-projecting nRE neurons consistently received substantially less prefrontal input than PFC-projecting nRE neurons (Fig. 2F; Welch’s two-sided t-test: t(4.73) = 4.58, P = 0.007). Thus, although nRE receives dense input from medial prefrontal cortex, these afferents largely avoid hippocampal-projecting neurons, with prominent input to these neurons coming instead from hypothalamus (Figure S4).

### PFC has limited functional connectivity with HPC-projecting nRE neurons

We next asked whether this anatomical segregation was reflected in functional connectivity. To directly compare prefrontal input onto the two projection-defined neuronal populations, we expressed channelrhodopsin-2 (ChR2) in medial prefrontal cortex and retrogradely labelled PFC- and HPC-projecting nRE neurons with fluorescent beads. Acute slices containing nRE were then used to perform sequential whole-cell recordings from neighbouring PFC- and HPC-projecting neurons while optogenetically stimulating the same population of prefrontal axons (Fig. 3A). Because both neurons were recorded within the same field of view, each pair experienced identical viral expression, slice conditions and optical stimulation, allowing a direct comparison of relative prefrontal synaptic input onto the two projection-defined populations.

**Figure 3.**
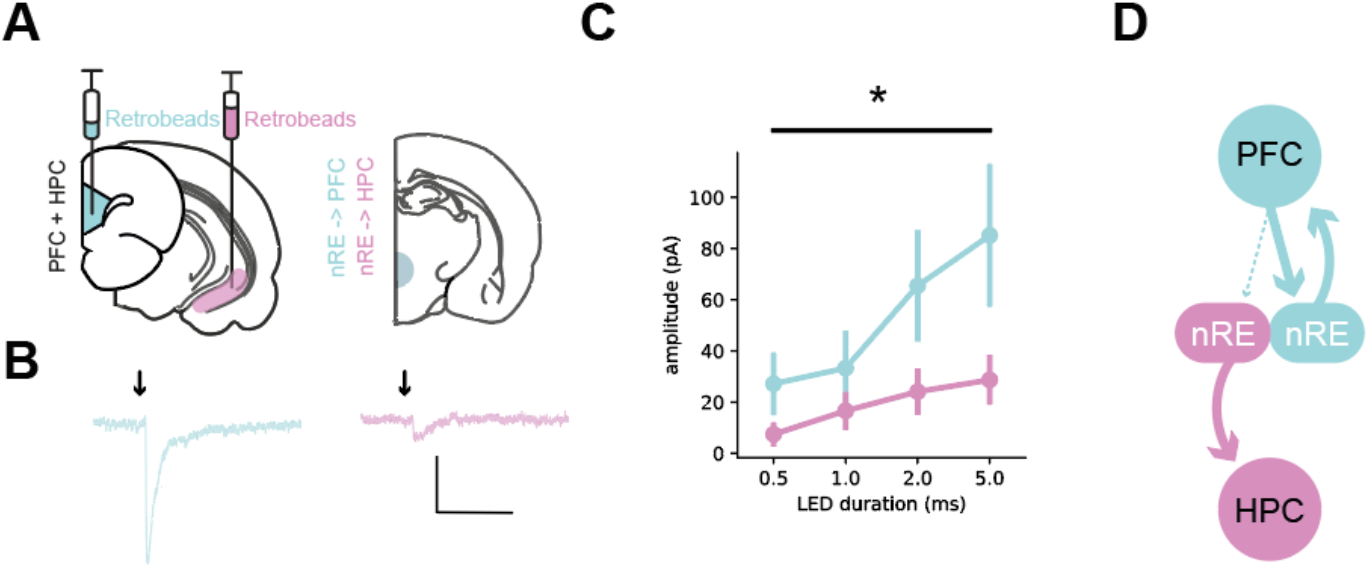
Prefrontal input exhibits weak functional connectivity with hippocampal-projecting nucleus reuniens neurons. (A) Experimental strategy for projection-specific circuit mapping. Channelrhodopsin-2 (ChR2) was expressed in medial prefrontal cortex (PFC), while retrobeads were injected into PFC (blue) and hippocampal formation (HPC; purple) to identify projection-defined nRE neurons. Acute brain slices were used for paired whole-cell recordings from neighbouring PFC- and HPC-projecting neurons during optogenetic stimulation of prefrontal afferents. (B) Representative light-evoked excitatory postsynaptic currents from PFC-projecting (blue) and HPC-projecting (purple) nRE neurons. Scale bar 50 pA, 100 ms. (C) Quantification of light-evoked synaptic responses across increasing light pulse durations (0.5–5 ms) in PFC- and HPC-projecting nRE neurons. (D) Summary schematic illustrating preferential functional connectivity of prefrontal afferents onto PFC-projecting rather than HPC-projecting nRE neurons.

Consistent with the tracing experiments, optical stimulation evoked robust excitatory currents in PFC-projecting neurons but much smaller, and frequently undetectable, responses in neighbouring HPC-projecting neurons (Fig. 3B,C). The difference became more pronounced with increasing light pulse duration (Fig. 3C, hierarchical linear mixed-effects model with cells nested within recording day; projection x intensity interaction, β = −36.8 ± 15.1, z = −2.44, P = 0.015). This concordance between anatomical and physiological measurements argues against technical explanations based on viral tropism or labelling efficiency.

Together, these anatomical and physiological data provide little support for the prevailing model in which medial prefrontal cortex communicates with hippocampus through a direct PFC–nRE–HPC relay. Instead, prefrontal cortex preferentially targets a distinct population of PFC-projecting nRE neurons, while hippocampal-projecting neurons receive remarkably little direct prefrontal input.

## DISCUSSION

Communication between prefrontal cortex (PFC) and hippocampus (HPC) is essential for flexible behaviour, working memory and episodic memory, and the thalamic nucleus reuniens (nRE) has long been proposed to provide the principal anatomical substrate for this interaction^9,31,32,42^. The prevailing model suggests that nRE functions as a relay, receiving information from PFC and transmitting it directly to HPC^9,31^. Here, by combining whole-brain anatomical mapping, projection-specific circuit tracing and functional connectivity measurements, we identify a circuit organisation that is inconsistent with this simple relay model. Although nRE is extensively and reciprocally connected with both PFC and HPC, prefrontal afferents exert little direct influence over hippocampal-projecting nRE neurons. Instead, these neurons receive highly selective input from hypothalamic regions, suggesting an alternative anatomical route through which prefrontal cortex may influence hippocampus^43–45^.

Our whole-brain mapping confirms and extends previous anatomical studies identifying nRE as a major point of interaction between PFC and HPC. Quantifying both afferent and efferent connectivity within a common anatomical framework demonstrates that nRE forms extensive reciprocal connections with both structures, reinforcing its central position within forebrain networks^31,32^. However, these anatomical relationships alone do not determine how information flows through the circuit. The widely accepted relay model implicitly assumes that prefrontal inputs converge onto the same nRE neurons that project to hippocampus.

We provide several independent observations that do not support this assumption. First, dual retrograde tracing showed that PFC- and HPC-projecting neurons comprise largely distinct projection-defined populations. Second, monosynaptic rabies tracing demonstrated that medial PFC contributes little direct input to hippocampal-projecting neurons despite providing robust input to neighbouring PFC-projecting neurons. Finally, optogenetic circuit mapping confirmed that functional excitatory input from PFC onto hippocampal-projecting neurons is weak or absent. The agreement between these independent anatomical and physiological approaches argues that the observed segregation reflects the underlying circuit organisation rather than limitations of any individual technique.

Together, these findings suggest that communication through nRE is organised through parallel projection-defined circuits rather than a common relay. Rather than acting as a passive conduit between PFC and HPC, distinct populations of nRE neurons receive different long-range inputs and are therefore likely to participate in different aspects of forebrain computation. Such an organisation is consistent with a growing appreciation that higher-order thalamic nuclei actively integrate and transform information, rather than simply transmitting it between connected regions^31,41,46–49^.

Our findings also fit with emerging evidence that communication through this circuit is organised to limit direct recurrent excitation between prefrontal cortex and hippocampus. We previously showed that nRE terminals within hippocampus largely avoid prefrontal-projecting hippocampal neurons^50^, reducing the potential for direct reciprocal excitation between these structures. Additionally, it has been shown that prefrontal-projecting hippocampal neurons do not collateralise to nRE^27^. The present study extends this organisational principle upstream, showing that hippocampal-projecting nRE neurons themselves receive remarkably little direct prefrontal input. Together, these observations suggest that communication between PFC and HPC is not mediated through a simple excitatory loop, but instead is subject to additional circuit-level control. One consequence of such an arrangement may be to reduce direct recurrent excitation within the PFC-HPC circuit, thereby promoting network stability. Consistent with this interpretation, recent work has shown that nRE input to CA1 and dorsal subiculum preferentially recruits local interneurons^51^, further limiting direct excitation within the circuit. These observations therefore raise the question of what afferent pathways instead provide the principal drive to hippocampal-projecting nRE neurons.

One particularly striking feature of the present study is the highly selective input from posterior and lateral hypothalamic regions onto hippocampal-projecting nRE neurons. These regions receive substantial prefrontal innervation^43^, suggesting a potential indirect pathway through which prefrontal cortex could influence hippocampal-projecting nRE neurons^45^. Although the functional significance of this pathway remains to be established, it provides a plausible anatomical substrate through which motivational and internal state signals carried by hypothalamus could influence hippocampal processing^43,52,53^.

Additional alternative routes exist, with prefrontal cortex projecting extensively to entorhinal and other parahippocampal regions^54^, which provide major cortical input to hippocampus^50,51^, as well as to amygdala and numerous other subcortical structures^55^ that also influence hippocampal processing^56^. These pathways may account for many effects previously attributed to direct prefrontal recruitment of hippocampal-projecting nRE neurons^31,36,37,57^. Future experiments that selectively compare these parallel pathways will be required to determine their relative contributions during behaviour. More generally, our results demonstrate that rather than principally by direct recruitment of hippocampal-projecting nRE neurons, communication between PFC and HPC through nRE may be predominantly mediated by intermediate subcortical circuits.

More broadly, our findings suggest that understanding nRE function will require moving beyond viewing it as a simple bridge between cortex and hippocampus. Instead, determining how distinct projection-defined neuronal populations integrate their different afferent inputs, and how these parallel circuits interact during behaviour, will be essential for understanding the contribution of nRE to cognition and memory. More generally, our findings suggest caution when interpreting the behavioural consequences of nRE manipulations as evidence for direct prefrontal recruitment of hippocampus.

## METHODS

### RESOURCE AVAILABILITY

#### Lead contact

Further information and requests for resources should be directed to and will be fulfilled by the Lead Contact, Andrew F. MacAskill.

#### Materials availability

This study did not generate unique reagents.

#### Data and code availability

The datasets and custom code generated during this study will be made publicly available upon publication. In the meantime they are available from the Lead Contact upon reasonable request.

## EXPERIMENTAL MODEL AND SUBJECT DETAILS

### Animals

All experiments were performed in adult male C57BL/6J mice (Charles River, UK) aged at least 6 weeks. Mice were housed under a 12 h light-dark cycle with food and water available ad libitum except where noted. All procedures were approved by the UK Home Office under the Animals (Scientific Procedures) Act and complied with institutional ethical guidelines.

## ANATOMY

### Stereotaxic surgery

Stereotaxic surgeries were performed as previously described^23,50,56^. Mice were anaesthetised with isoflurane (4% induction, 1.5–2% maintenance) and secured in a stereotaxic frame (Kopf Instruments). Following a midline scalp incision, craniotomies were made over the target coordinates and injections were performed using pulled glass micropipettes connected to a Nanoject II injector (Drummond Scientific). Viral vectors and tracers were delivered in 27.6 nL aliquots every 10 s, after which the pipette was left in place for 1–5 min before slow withdrawal to minimise reflux. Injections targeting nucleus reuniens were performed using a 10° approach angle to avoid the superior sagittal sinus. Following surgery, the incision was sutured and animals recovered on a heating pad. Carprofen (Rimadyl; 5 mg kg^−1^, s.c.) was administered peri-operatively, with additional carprofen (0.05 mg mL^−1^) provided in the drinking water for 48 h post-operatively. Animals were allowed to recover for at least 2 weeks before subsequent procedures unless otherwise stated.

Injections were targeted to infralimbic medial prefrontal cortex (AP +1.6, ML ±0.2, DV −2.5), ventral hippocampus (CA1: AP −3.8, ML ±3.1, DV −4.1 to −4.3, subiculum: AP: −3.9, ML ±3.1, DV −2.0 to −2.1), and nucleus reuniens (AP −0.7, ML ±0.9, DV −4.5), with nRE injections performed using a 10° approach angle to avoid the superior sagittal sinus. All coordinates are relative to bregma.

### Retrograde CTB tracing

To map long-range inputs, Alexa Fluor 555- or 647-conjugated cholera toxin B (CTB; 250 nL, 1 mg mL^−1)^ was injected into nucleus reuniens, medial prefrontal cortex or hippocampus. For experiments examining collateralisation of prefrontal and hippocampal projections, spectrally distinct CTB tracers were injected simultaneously into prefrontal cortex and hippocampus. After a minimum of 14 days, mice were perfused and brains sectioned coronally (60 μm). Every second section spanning the rostrocaudal extent of the brain was collected and imaged using a Zeiss Axioscan.

### Monosynaptic rabies tracing

Projection-defined monosynaptic rabies tracing was performed using a Cre-dependent helper strategy^50,58^. AAV1-hSyn-Cre-WPRE-hGH (Addgene #105553) was injected into medial prefrontal cortex or hippocampus together with AAV2/1-synP-FLEX-split-TVA-EGFP-B19G (Addgene #52473 and Charité BA-96) into nucleus reuniens. After at least two weeks, EnvA-pseudotyped glycoprotein-deleted rabies virus expressing mCherry (BRV-envA-1d-mCherry) was injected into nucleus reuniens. Animals were perfused seven days later. Brains were imaged using serial two-photon tomography.

### Whole-brain image registration and cell detection

CTB tracing and monosynaptic rabies tracing datasets were processed using independent image analysis pipelines optimised for their respective imaging modalities before being analysed within a common anatomical framework.

For CTB tracing experiments, every second 60 μm coronal section spanning the rostrocaudal extent of the brain was imaged using a Zeiss Axioscan. Coronal sections were registered to the Allen Mouse Brain Atlas using the WholeBrain framework^59^. Initial anteroposterior coordinates were assigned manually to landmark sections and interpolated across the remaining sections before automated registration, which was refined by manual adjustment of anatomical landmarks where necessary. Retrogradely labelled neurons were detected using wavelet-based multiresolution decomposition. Automated segmentations were manually curated in Napari to remove false-positive detections arising from uneven illumination or tissue artefacts. For dual-tracer experiments, cell detection was performed independently for each fluorescence channel, and dual-labelled neurons were identified by manual confirmation of colocalisation in the underlying images.

For monosynaptic rabies tracing experiments, serial two-photon tomography datasets were registered to the Allen Common Coordinate Framework (CCFv3) using brainreg^60,61^. Labelled neurons were detected using cellfinder^62^ with a pretrained convolutional neural network and visualised using brainrender^63^. Brain regions consistently associated with the injection site or injection tract were excluded from subsequent analyses. Brains were then manually curated to correct false-negative and remove false-positive detections arising from uneven illumination or tissue artefacts. Because GFP expression from helper virus-expressing starter cells was imaged under acquisition conditions optimised for detection of rabies-labelled neurons, starter cells could not be reliably segmented from the tomography datasets. Consequently, analyses were based on verification of injection location rather than quantitative assessment of starter cell number or distribution.

Coordinates obtained from either WholeBrain or brainreg were transformed into a common Allen Common Coordinate Framework, allowing anatomical identities to be assigned using the Allen Brain Atlas hierarchy through the Allen SDK. Brain regions were grouped according to predefined Allen anatomical ontologies to permit quantitative comparisons between CTB and rabies datasets despite their independent acquisition and registration pipelines.

### Quantification of anatomical connectivity

For each animal, labelled neurons assigned to each anatomical region were expressed as a proportion of the total number of labelled neurons to account for differences in overall labelling efficiency between experiments. Anatomical quantification was performed on biological replicates (individual animals) rather than individual cells. Injection sites were verified manually with reference to the Allen Mouse Brain Atlas, and animals with inaccurate targeting were excluded from further analysis. Statistical comparisons were performed on regional proportions across animals.

## ELECTROPHYSIOLOGY

### Slice preparation

Acute coronal brain slices (300 μm) containing nucleus reuniens were prepared from adult mice using approaches described previously^23,50,56^, following at least 3 weeks of viral expression. Mice were deeply anaesthetised with ketamine/xylazine and transcardially perfused with an ice-cold sucrose-based cutting solution containing (in mM): 190 sucrose, 25 glucose, 10 NaCl, 25 NaHCO_3_, 1.2 NaH_2_PO_4_, 2.5 KCl, 1 sodium ascorbate, 2 sodium pyruvate, 7 MgCl_2_ and 0.5 CaCl_2_, continuously bubbled with 95% O_2_ and 5% CO_2_. Brains were rapidly removed and 300 μm coronal slices prepared using a vibrating microtome. Slices were transferred to artificial cerebrospinal fluid (aCSF) containing (in mM): 125 NaCl, 22.5 glucose, 25 NaHCO_3_, 1.25 NaH_2_PO_4_, 2.5 KCl, 1 sodium ascorbate, 3 sodium pyruvate, 1 MgCl_2_ and 2 CaCl_2_, equilibrated with 95% O_2_/5% CO_2_, incubated at 35 °C for 30 min and subsequently maintained at room temperature until recording.

### Whole-cell recordings

Whole-cell voltage-clamp recordings were obtained from nucleus reuniens neurons retrogradely labelled by injection of red or green RetroBeads (Lumofluor) into medial prefrontal cortex or ventral hippocampus. Projection-defined neurons were identified by fluorescence and targeted under Dodt contrast optics. Borosilicate recording pipettes (2–4 MΩ) were filled with a Cs-gluconate-based internal solution containing (in mM): 135 Cs-gluconate, 10 HEPES, 7 KCl, 10 sodium phosphocreatine, 4 MgATP, 0.4 NaGTP, 10 TEA and 2 QX-314. Excitatory postsynaptic currents were recorded at a holding potential of −70 mV in the presence of 10 μM gabazine to isolate glutamatergic transmission. Signals were acquired using a Multiclamp 700B amplifier, filtered at 4 kHz and digitised at 10 kHz.

### Optogenetic stimulation

To selectively stimulate prefrontal cortical afferents, AAV1-hSyn-hChR2(H134R)-EYFP was injected into infralimbic medial prefrontal cortex. ChR2-expressing axons within nucleus reuniens were stimulated using brief pulses of blue light (473 nm LED; CoolLED pE-4000) delivered through a 40× objective. Light pulses of increasing duration (0.5, 1, 2 and 5 ms) were presented while recording from hippocampal-projecting and prefrontal-projecting nucleus reuniens neurons. Light intensity was maintained constant across recordings (4–7 mW measured at the back aperture of the objective). Peak evoked EPSC amplitudes were quantified offline and compared between projection-defined neuronal populations.

## STATISTICAL ANALYSIS

Statistical analyses were performed in Python using Pingouin and statsmodels. Whole-brain anatomical tracing data are presented descriptively as the proportion of labelled neurons assigned to each anatomical region for each animal. Differences in prefrontal cortical input to hippocampal- and prefrontal-projecting nucleus reuniens neurons were assessed using a Welch’s two-sided independent-samples t-test. Electrophysiological recordings were analysed using hierarchical linear mixed-effects models with projection target, stimulus duration and their interaction included as fixed effects, and cells nested within each recording day as random effects. Exact sample sizes, test statistics and p values are reported in the text.

## ACKNOWLEDGEMENTS

We thank members of the MacAskill laboratory for helpful comments on the manuscript. A.F.M. was supported by a UKRI Frontier Research Fellowship (grant number EP/Y034724/1). J.S. was supported by the Wellcome Trust 4-year PhD in Neuroscience at UCL (grant number 222292/Z/20/Z).

## AUTHOR CONTRIBUTIONS

Conceptualization, Methodology, Investigation, Formal Analysis, Writing – Original Draft, Writing – Review & Editing, J.S., K.V. and A.F.M.; Funding Acquisition, Supervision, A.F.M.

## DECLARATION OF INTERESTS

The authors declare no competing interests.

## INCLUSION AND DIVERSITY

We support inclusive, diverse, and equitable conduct of research.

---

**Figure S1.**
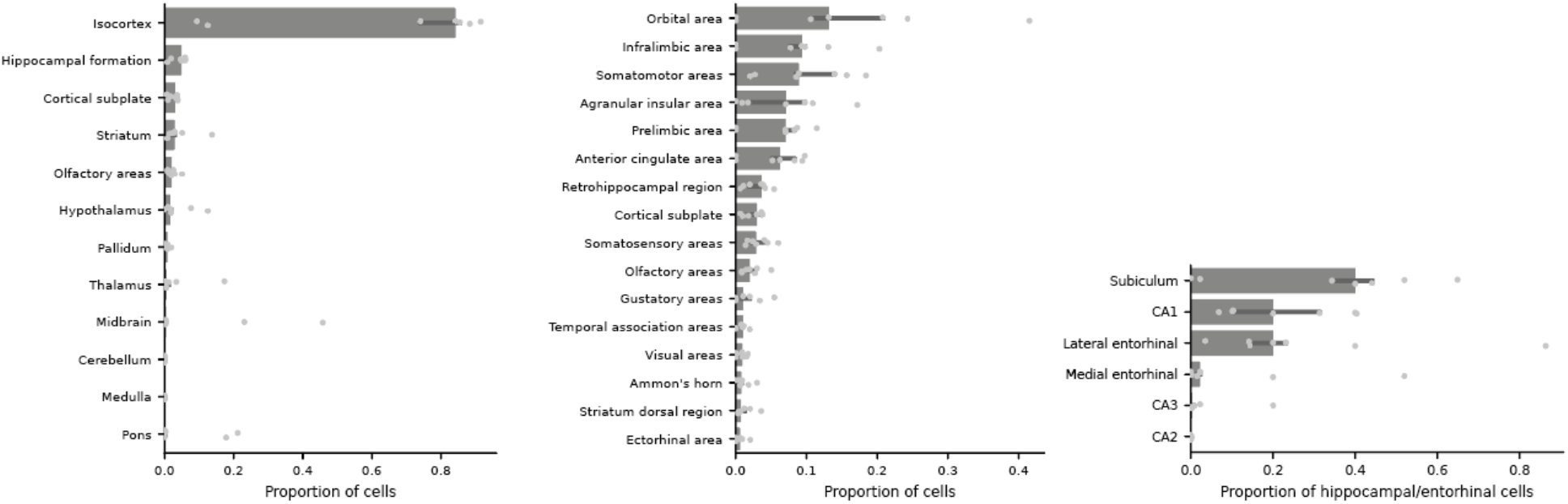
Quantification of brain-wide afferent inputs to nucleus reuniens. (A) Proportion of CTB-labelled neurons across major brain divisions following injections into nucleus reuniens. (B) Proportion of cortical inputs across individual cortical subregions. (C) Proportion of hippocampal formation inputs across hippocampal subfields.

**Figure S2.**
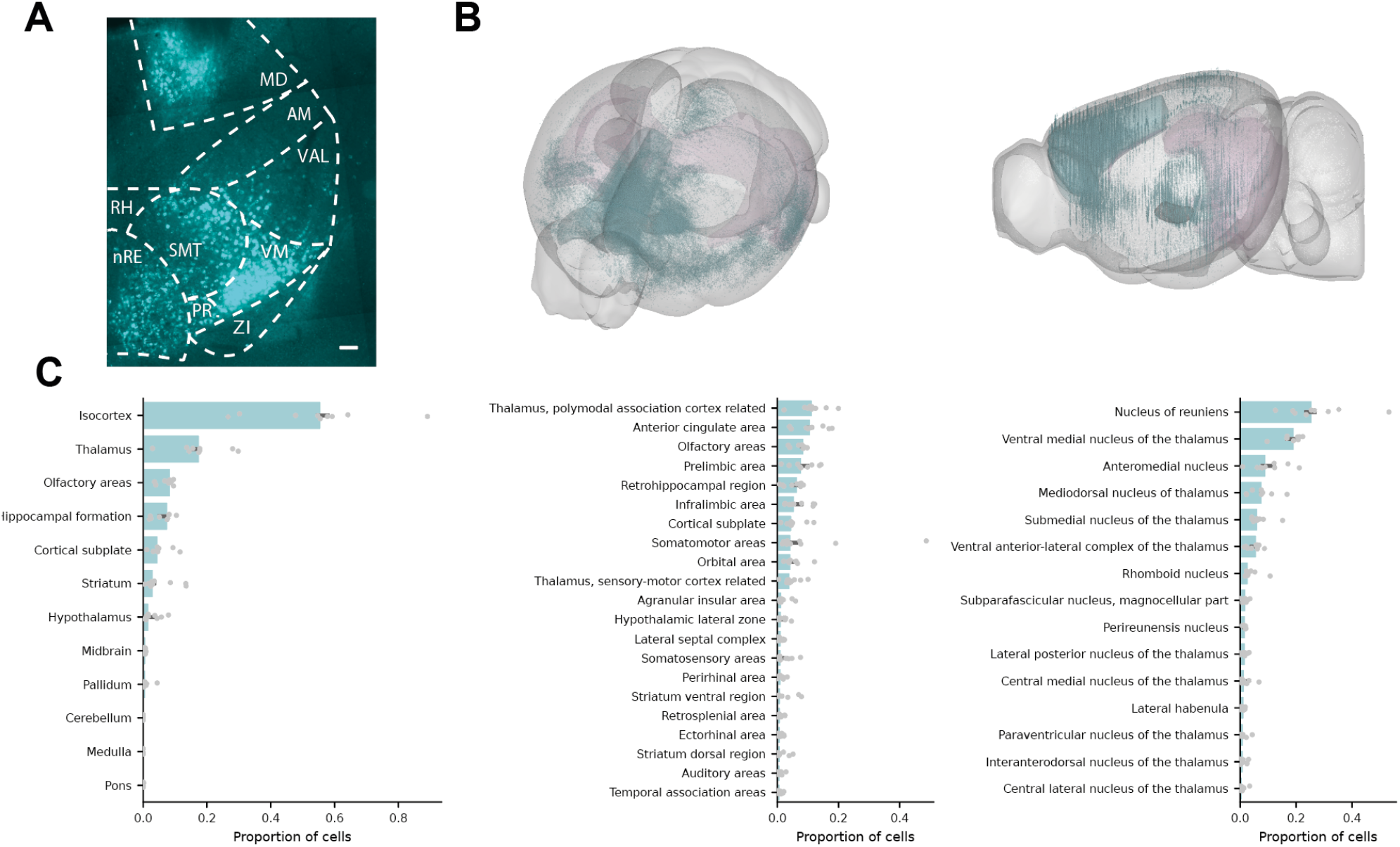
Brain-wide quantification of afferent inputs to prefrontal cortex. (A) Representative coronal section showing CTB-labelled neurons in thalamus following CTB injection into prefrontal cortex. (B) Whole-brain distribution of labelled neurons registered to the Allen Common Coordinate Framework. (C) Quantification of labelled neurons. Left, proportion of labelled neurons across major brain divisions. Middle, proportion of labelled neurons across individual brain regions. Right, proportion of labelled thalamic neurons across thalamic subregions.

**Figure S3.**
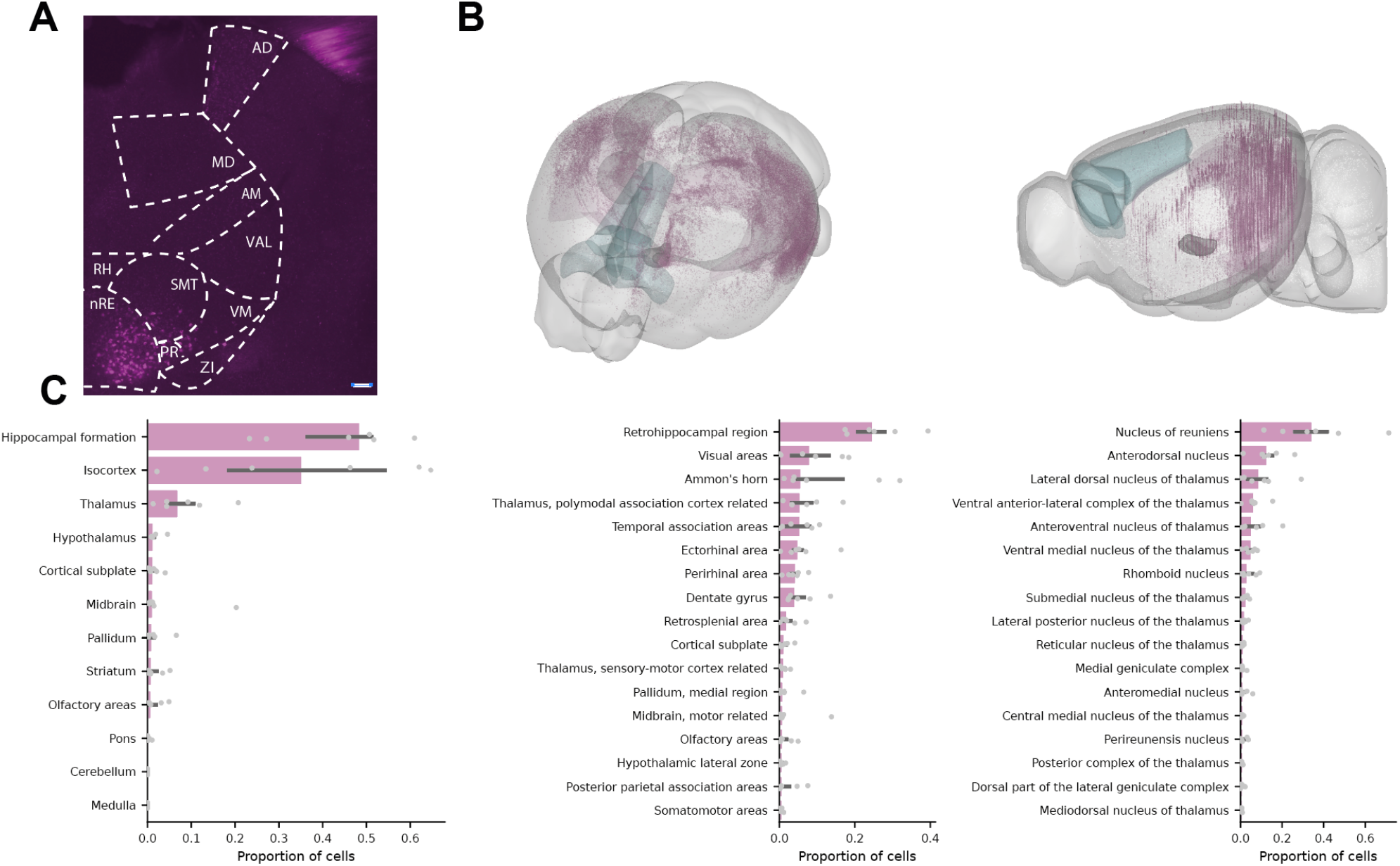
Brain-wide quantification of afferent inputs to hippocampus. (A) Representative coronal section showing CTB-labelled neurons following CTB injection into hippocampus. (B) Whole-brain distribution of labelled neurons registered to the Allen Common Coordinate Framework. (C) Quantification of labelled neurons. Left, proportion of labelled neurons across major brain divisions. Middle, proportion of labelled neurons across individual brain regions. Right, proportion of labelled thalamic neurons across thalamic subregions.

**Figure S4.**
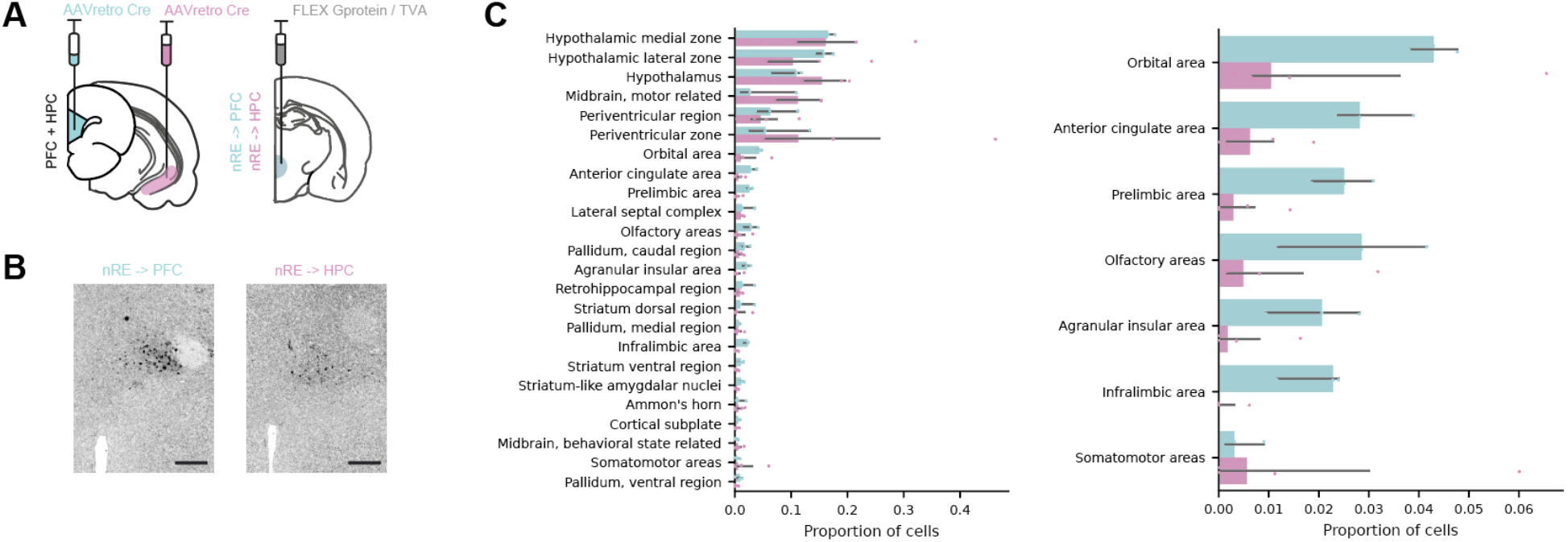
Projection-specific monosynaptic rabies tracing of prefrontal cortex- and hippocampal-projecting nucleus reuniens neurons. (A) Schematic of the projection-specific monosynaptic rabies tracing strategy using retrograde AAV-Cre, Cre-dependent TVA/G helper viruses, and EnvA-pseudotyped glycoprotein-deleted rabies virus. (B) Representative images of starter cells in nucleus reuniens. Images were acquired using settings optimised for detection of the red rabies fluorophore, resulting in comparatively weak helper virus fluorescence. (C) Quantification of monosynaptic inputs to prefrontal cortex-projecting (blue) and hippocampal-projecting (purple) nucleus reuniens neurons. Left, proportion of labelled neurons across major brain divisions. Right, proportion of labelled neurons across prefrontal cortical subregions.

## Notes

### Competing Interest Statement

The authors have declared no competing interest.

## REFERENCES

1. Totty, M.S., Tuna, T., Ramanathan, K.R., Jin, J., Peters, S.E., and Maren, S. (2023). Thalamic nucleus reuniens coordinates prefrontal-hippocampal synchrony to suppress extinguished fear. Nature Communications 14. 10.1038/s41467-023-42315-1.

2. de Mooij-van Malsen, J.G., Röhrdanz, N., Buschhoff, A.S., Schiffelholz, T., Sigurdsson, T., and Wulff, P. (2023). Task-specific oscillatory synchronization of prefrontal cortex, nucleus reuniens, and hippocampus during working memory. iScience 26. 10.1016/j.isci.2023.107532.

3. Griffin, A.L. (2021). The nucleus reuniens orchestrates prefrontal-hippocampal synchrony during spatial working memory. Neuroscience and Biobehavioral Reviews 128, 415–420. 10.1016/j.neubiorev.2021.05.033.

4. Jayachandran, M., Linley, S.B., Schlecht, M., Mahler, S.V., Vertes, R.P., and Allen, T.A. (2019). Prefrontal Pathways Provide Top-Down Control of Memory for Sequences of Events. Cell Reports 28, 640–654.e6. 10.1016/j.celrep.2019.06.053.

5. Mölle, M., and Born, J. (2011). Slow oscillations orchestrating fast oscillations and memory consolidation. In Progress in Brain Research (Elsevier B.V.), pp. 93–110. 10.1016/B978-0-444-53839-0.00007-7.

6. Wirt, R.A., and Hyman, J.M. (2017). Integrating spatial working memory and remote memory: Interactions between the medial prefrontal cortex and hippocampus. Brain Sciences 7. 10.3390/brainsci7040043.

7. Eichenbaum, H. (2017). Prefrontal–hippocampal interactions in episodic memory. Nat Rev Neurosci 18, 547–558. 10.1038/nrn.2017.74.

8. Wikenheiser, A.M., and Schoenbaum, G. (2016). Over the river, through the woods: cognitive maps in the hippocampus and orbitofrontal cortex. Nat Rev Neurosci 17, 513–523. 10.1038/nrn.2016.56.

9. Jin, J., and Maren, S. (2015). Prefrontal-Hippocampal Interactions in Memory and Emotion. Frontiers Syst Neurosci 9, 170. 10.3389/fnsys.2015.00170.

10. Passlack, J., and MacAskill, A.F. (2026). Contextual inference through flexible integration of environmental features and behavioural outcomes. PLOS Computational Biology 22, e1014093. 10.1371/journal.pcbi.1014093.

11. Guise, K.G., and Shapiro, M.L. (2017). Medial Prefrontal Cortex Reduces Memory Interference by Modifying Hippocampal Encoding. Neuron 94, 183–192. 10.1016/j.neuron.2017.03.011.

12. Hallock, H.L., Wang, A., and Griffin, A.L. (2016). Ventral midline thalamus is critical for hippocampal–prefrontal synchrony and spatial working memory. Journal of Neuroscience 36, 8372–8389. 10.1523/JNEUROSCI.0991-16.2016.

13. Ito, H.T., Zhang, S.J., Witter, M.P., Moser, E.I., and Moser, M.B. (2015). A prefrontal-thalamo-hippocampal circuit for goal-directed spatial navigation. Nature 522, 50–55. 10.1038/nature14396.

14. Siapas, A.G., Lubenov, E.V., and Wilson, M.A. (2005). Prefrontal phase locking to hippocampal theta oscillations. Neuron 46, 141–151. 10.1016/j.neuron.2005.02.028.

15. Fujisawa, S., and Buzsáki, G. (2011). A 4 Hz Oscillation Adaptively Synchronizes Prefrontal, VTA, and Hippocampal Activities. Neuron 72, 153–165. 10.1016/j.neuron.2011.08.018.

16. Spellman, T., Rigotti, M., Ahmari, S.E., Fusi, S., Gogos, J.A., and Gordon, J.A. (2015). Hippocampal-prefrontal input supports spatial encoding in working memory. Nature 522, 309–314. 10.1038/nature14445.

17. Ferraris, M., Cassel, J.C., Pereira de Vasconcelos, A., Stephan, A., and Quilichini, P.P. (2021). The nucleus reuniens, a thalamic relay for cortico-hippocampal interaction in recent and remote memory consolidation. Neuroscience and Biobehavioral Reviews 125, 339–354. 10.1016/j.neubiorev.2021.02.025.

18. Bloodgood, D.W., Sugam, J.A., Holmes, A., and Kash, T.L. (2018). Fear extinction requires infralimbic cortex projections to the basolateral amygdala. Translational Psychiatry 8. 10.1038/s41398-018-0106-x.

19. Zhao, X., Hsu, C.L., and Spruston, N. (2022). Rapid synaptic plasticity contributes to a learned conjunctive code of position and choice-related information in the hippocampus. Neuron 110, 96–108. 10.1016/j.neuron.2021.10.003.

20. Frankland, P.W., Filipkowski, R.K., Cestari, V., McDonald, R.J., and Silva, A.J. (1998). The dorsal hippocampus is essential for context discrimination but not for contextual conditioning. Behavioral Neuroscience 112, 863–874. 10.1037/0735-7044.112.4.863.

21. Ainge, J.A., Van Der Meer, M.A.A., Langston, R.F., and Wood, E.R. (2007). Exploring the Role of Context-Dependent Hippocampal Activity in Spatial Alternation Behavior. Hippocampus 17, 988–1002. 10.1002/hipo.

22. Hoover, W.B., and Vertes, R.P. (2007). Anatomical analysis of afferent projections to the medial prefrontal cortex in the rat. Brain Structure and Function 212, 149–179. 10.1007/s00429-007-0150-4.

23. Sánchez-Bellot, C., and MacAskill, A.F. (2022). Push-pull regulation of exploratory behavior by two opposing hippocampal to prefrontal cortex pathways. Nature Communications 13. 10.1038/s41467-022-27977-7.

24. Sesack, S.R., Deutch, A.Y., Roth, R.H., and Bunney, B.S. (1989). Topographical organization of the efferent projections of the medial prefrontal cortex in the rat: An anterograde tract-tracing study with Phaseolus vulgaris leucoagglutinin. Journal of Comparative Neurology 290, 213–242. 10.1002/cne.902900205.

25. Vertes, R.P. (2002). Analysis of projections from the medial prefrontal cortex to the thalamus in the rat, with emphasis on nucleus reuniens. Journal of Comparative Neurology, 163–187. 10.1002/cne.10083.

26. McKenna, J.T., and Vertes, R.P. (2004). Afferent projections to nucleus reuniens of the thalamus. Journal of Comparative Neurology 480, 115–142. 10.1002/cne.20342.

27. Varela, C., Kumar, S., Yang, J.Y., and Wilson, M.A. (2014). Anatomical substrates for direct interactions between hippocampus, medial prefrontal cortex, and the thalamic nucleus reuniens. Brain Structure and Function 219, 911–929. 10.1007/s00429-013-0543-5.

28. Dolleman-van der Weel, M.J., Lopes da Silva, F.H., and Witter, M.P. (2017). Interaction of nucleus reuniens and entorhinal cortex projections in hippocampal field CA1 of the rat. Brain Structure and Function 222, 2421–2438. 10.1007/s00429-016-1350-6.

29. Vertes, R.P., Hoover, W.B., Do Valle, A.C., Sherman, A., and Rodriguez, J.J. (2006). Efferent projections of reuniens and rhomboid nuclei of the thalamus in the rat. Journal of Comparative Neurology 499, 768–796. 10.1002/cne.21135.

30. Wouterlood, F.G., Saldana, E., and Witter, M.P. (1990). Projection from the nucleus reuniens thalami to the hippocampal region: Light and electron microscopic tracing study in the rat with the anterograde tracer Phaseolus vulgaris-leucoagglutinin. Journal of Comparative Neurology 296, 179–203. 10.1002/cne.902960202.

31. Dolleman-Van Der Weel, M.J., Griffin, A.L., Ito, H.T., Shapiro, M.L., Witter, M.P., Vertes, R.P., and Allen, T.A. (2019). The nucleus reuniens of the thalamus sits at the nexus of a hippocampus and medial prefrontal cortex circuit enabling memory and behavior. Learning and Memory 26, 191–205. 10.1101/lm.048389.118.

32. Tuna, T., and Maren, S. (2026). The thalamic nucleus reuniens orchestrates prefrontal-hippocampal synchrony in memory, emotion, and disease. Frontiers in Behavioral Neuroscience 20. 10.3389/fnbeh.2026.1885971.

33. Vertes, R.P., Hoover, W.B., Szigeti-Buck, K., and Leranth, C. (2007). Nucleus reuniens of the midline thalamus: Link between the medial prefrontal cortex and the hippocampus. Brain Research Bulletin 71, 601–609. 10.1016/j.brainresbull.2006.12.002.

34. Cassel, J.C., Ferraris, M., Quilichini, P., Cholvin, T., Boch, L., Stephan, A., and Pereira de Vasconcelos, A. (2021). The reuniens and rhomboid nuclei of the thalamus: A crossroads for cognition-relevant information processing? Neuroscience and Biobehavioral Reviews 126, 338–360. 10.1016/j.neubiorev.2021.03.023.

35. Eleore, L., López-Ramos, J.C., Guerra-Narbona, R., and Delgado-García, J.M. (2011). Role of reuniens nucleus projections to the medial prefrontal cortex and to the hippocampal pyramidal CA1 area in associative learning. PLoS ONE 6. 10.1371/journal.pone.0023538.

36. Xu, W., and Südhof, T.C. (2013). A neural circuit for memory specificity and generalization. Science 339, 1290–1295. 10.1126/science.1229534.

37. Zimmerman, E.C., and Grace, A.A. (2018). Prefrontal cortex modulates firing pattern in the nucleus reuniens of the midline thalamus via distinct corticothalamic pathways. European Journal of Neuroscience 48, 3255–3272. 10.1111/ejn.14111.

38. Lara-Vásquez, A., Espinosa, N., Durán, E., Stockle, M., and Fuentealba, P. (2016). Midline thalamic neurons are differentially engaged during hippocampus network oscillations. Scientific Reports 6. 10.1038/srep29807.

39. Steriade, M., and Deschenes, M. (1984). The Thalamus As a Neuronal Oscillator.

40. Cruz, K.G., Leow, Y.N., Le, N.M., Adam, E., Huda, R., and Sur, M. (2023). Cortical-subcortical interactions in goal-directed behavior. Physiological Reviews 103, 347–389. 10.1152/physrev.00048.2021.

41. Vantomme, G., Devienne, G., Hull, J.M., and Huguenard, J.R. (2025). The reuniens thalamus recruits recurrent excitation in the medial prefrontal cortex. Proceedings of the National Academy of Sciences 122, e2500321122. 10.1073/pnas.2500321122.

42. Eichenbaum, H. (2017). Prefrontal-hippocampal interactions in episodic memory. Nature Reviews Neuroscience 18, 547–558. 10.1038/nrn.2017.74.

43. González, J.A., Iordanidou, P., Strom, M., Adamantidis, A., and Burdakov, D. (2016). Awake dynamics and brain-wide direct inputs of hypothalamic MCH and orexin networks. Nature Communications 7. 10.1038/ncomms11395.

44. Mohammad, H., Senol, E., Graf, M., Lee, C.Y., Li, Q., Liu, Q., Yeo, X.Y., Wang, M., Laskaratos, A., Xu, F., et al. (2021). A neural circuit for excessive feeding driven by environmental context in mice. Nature Neuroscience 24, 1132–1141. 10.1038/s41593-021-00875-9.

45. Supiot, L., Benschop, A., Haak, N., Wolterink-Donselaar, I., Luijendijk, M., Adan, R., Poorthuis, R., and Meye, F. (2024). A prefrontal cortex-lateral hypothalamus circuit controls stress-driven food intake. bioRxiv. 10.1101/2024.05.02.592146.

46. Arber, S., and Costa, R.M. (2022). Networking brainstem and basal ganglia circuits for movement. Nature Reviews Neuroscience 23. 10.1038/s41583-022-00581-w.

47. Schmitt, L.I., Wimmer, R.D., Nakajima, M., Happ, M., Mofakham, S., and Halassa, M.M. (2017). Thalamic amplification of cortical connectivity sustains attentional control. Nature 545, 219–223. 10.1038/nature22073.

48. Hummos, A., Wang, B.A., Drammis, S., Halassa, M.M., and Pleger, B. (2022). Thalamic regulation of frontal interactions in human cognitive flexibility. PLoS Computational Biology 18. 10.1371/journal.pcbi.1010500.

49. Rikhye, R.V., Wimmer, R.D., and Halassa, M.M. (2026). Toward an Integrative Theory of Thalamic Function. 125, 12–12. 10.1146/annurev-neuro-080317.

50. Wee, R.W.S., and MacAskill, A.F. (2020). Biased Connectivity of Brain-wide Inputs to Ventral Subiculum Output Neurons. Cell Reports 30, 3644-3654.e6. 10.1016/j.celrep.2020.02.093.

51. Andrianova, L., Banks, P.J., Booth, C.A., Brady, E.S., Margetts-Smith, G., Kohli, S., Cavanagh, J., Bashir, Z.I., McBain, C.J., and Craig, M.T. (2025). Thalamic nucleus reuniens preferentially targets inhibitory interneurons over pyramidal cells in hippocampal CA1 region. biorxiv. 10.1101/2021.09.30.462517.

52. Sakurai, T. (2014). The role of orexin in motivated behaviours. Nature Reviews Neuroscience 15, 719–731. 10.1038/nrn3837.

53. Sharpe, M.J. (2024). The cognitive (lateral) hypothalamus. Trends in Cognitive Sciences 28, 18–29. 10.1016/j.tics.2023.08.019.

54. Uylings, H.B.M., Groenewegen, H.J., and Kolb, B. (2003). Do rats have a prefrontal cortex? Behavioural Brain Research 146, 3–17. 10.1016/j.bbr.2003.09.028.

55. Gao, L., Liu, S., Gou, L., Hu, Y., Liu, Y., Deng, L., Ma, D., Wang, H., Yang, Q., Chen, Z., et al. (2022). Single-neuron projectome of mouse prefrontal cortex. Nature Neuroscience 25, 515–529. 10.1038/s41593-022-01041-5.

56. AlSubaie, R., Wee, R.W., Ritoux, A., Mishchanchuk, K., Passlack, J., Regester, D., and MacAskill, A.F. (2021). Control of parallel hippocampal output pathways by amygdalar long-range inhibition. Elife 10, e74758. 10.7554/elife.74758.

57. Cheung, H., Yu, T.Z., Yi, X., Wu, Y.J., Wang, Q., Gu, X., Xu, M., Cai, M., Wen, W., Li, X.N., et al. (2024). An ultra-short-acting benzodiazepine in thalamic nucleus reuniens undermines fear extinction via intermediation of hippocamposeptal circuits. Communications Biology 7. 10.1038/s42003-024-06417-w.

58. Beier, K.T., Steinberg, E.E., Deloach, K.E., Xie, S., Miyamichi, K., Schwarz, L., Gao, X.J., Kremer, E.J., Malenka, R.C., and Luo, L. (2015). Circuit Architecture of VTA Dopamine Neurons Revealed by Systematic Input-Output Mapping. Cell 162, 622–634. 10.1016/j.cell.2015.07.015.

59. Fürth, D., Vaissière, T., Tzortzi, O., Xuan, Y., Märtin, A., Lazaridis, I., Spigolon, G., Fisone, G., Tomer, R., Deisseroth, K., et al. (2018). An interactive framework for whole-brain maps at cellular resolution. Nature Neuroscience 21, 895–895. 10.1038/s41593-017-0058-0.

60. Claudi, F., Petrucco, L., Tyson, A., Branco, T., Margrie, T., and Portugues, R. (2020). BrainGlobe Atlas API: a common interface for neuroanatomical atlases. Journal of Open Source Software 5, 2668–2668. 10.21105/joss.02668.

61. Tyson, A.L., Vélez-Fort, M., Rousseau, C.V., Cossell, L., Tsitoura, C., Lenzi, S.C., Obenhaus, H.A., Claudi, F., Branco, T., and Margrie, T.W. (2022). Accurate determination of marker location within whole-brain microscopy images. Scientific Reports 12. 10.1038/s41598-021-04676-9.

62. Tyson, A.L., Rousseau, C.V., Niedworok, C.J., Keshavarzi, S., Tsitoura, C., Cossell, L., Strom, M., and Margrie, T.W. (2021). A deep learning algorithm for 3D cell detection in whole mouse brain image datasets. PLoS Computational Biology 17. 10.1371/journal.pcbi.1009074.

63. Claudi, F., Tyson, A.L., Petrucco, L., Margrie, T.W., Portugues, R., and Branco, T. (2021). Visualizing anatomically registered data with brainrender. eLife 10. 10.7554/eLife.65751.

